# Metagenomic discovery of *Candidatus* Parvarchaeales related lineages sheds light on the adaptation and diversification from neutral-thermal to acidic-mesothermal environments

**DOI:** 10.1101/2022.12.14.520523

**Authors:** Yang-Zhi Rao, Yu-Xian Li, Ze-Wei Li, Yan-Ni Qu, Yan-Ling Qi, Jian-Yu Jiao, Wen-Sheng Shu, Zheng-Shuang Hua, Wen-Jun Li

**Author notes:** Correspondence and requests for materials should be addressed to Z.S.H and W.J.L. These authors contributed equally to this work: Yang-Zhi Rao, Yu-Xian Li. **Authors’ e-mail address list:** Y.Z.R. Y.X.L. L.Z.W. Y.N.Q. Y.L.Q. J.Y.J. W.S.S. Z.S.H. W.J.L.

## Abstract

*Candidatus* Parvarchaeales, representing a DPANN archaeal group with limited metabolic potentials and reliance on hosts for their growth, were initially found in acid mine drainage (AMD). Due to the lack of representatives, however, their ecological roles and adaptation to extreme habitats such as AMD, as well as how they diverge across the lineage remain largely unexplored. By applying genome-resolved metagenomics, 28 *Parvarchaeales*-associated metagenome-assembled genomes (MAGs) representing two orders and five genera were recovered. Among them, we identified three new genera and proposed the names *Candidatus* Jingweiarchaeum, *Candidatus* Haiyanarchaeum, and *Candidatus* Rehaiarchaeum with the former two belonging to a new order *Candidatus* Jingweiarchaeales. Further analyses of metabolic potentials revealed substantial niche differentiation between Jingweiarchaeales and Parvarchaeales. Jingweiarchaeales may rely on fermentation, salvage pathways, partial glycolysis, and pentose phosphate pathway (PPP) for energy reservation, while the metabolic potentials of Parvarchaeales might be more versatile. Comparative genomic analyses suggested that Jingweiarchaeales are more favorable to habitats with higher temperatures and *Parvarchaeales* are better adapted to acidic environments. We further revealed that the thermal adaptation of these lineages especially for Haiyanarchaeum might rely on innate genomic features such as the usage of specific amino acids, genome streamlining, and hyperthermal featured genes such as *rgy*. Notably, the acidic adaptation of Parvarchaeales was possibly driven by horizontal gene transfer (HGT). Reconstruction of ancestral states demonstrated that both may originate from thermal and neutral environments and later spread to mesothermal and acidic environments. These evolutionary processes may also be accompanied by adaptation toward oxygen-rich environments via HGT.

**Importance:** *Candidatus* Parvarchaeales may represent a lineage uniquely distributed in extreme environments such as AMD and hot springs. However, little is known about the strategies and processes of how they adapted to these extreme environments. By the discovery of potential new order-level lineages - Jingweiarchaeales and in-depth comparative genomic analysis, we unveiled the functional differentiation of these lineages. Further, we show that the adaptation to high-temperature and acidic environments of these lineages was driven by different strategies, with the prior relying more on innate genomic characteristics and the latter more on the acquisition of genes associated with acid tolerance. Finally, by reconstruction of ancestral states of OGT and *pI*, we showed the potential evolutionary process of Parvarchaeales-related lineages with regard to the shift from a high-temperature environment of their common ancestors to low-temperature (potentially acidic) environments.

## Introduction

Archaea were among the earliest emerging lineages of the tree of life (Williams et al. 2020; Woese and Fox 1977). Diversely found in all sorts of environments, Archaea often predominate extreme environments such as hot springs, acid mine drainages (AMD), and hypersaline environments (Shu and Huang 2022). As a major archaeal lineage, DPANN archaea have uniquely small genome sizes with limited metabolic potentials and are thus considered to have a symbiotic lifestyle (Castelle et al. 2015; Chen et al. 2018; Huber et al. 2002; Li et al. 2021; Liu et al. 2018; Parks et al. 2017; Probst et al. 2018; Rinke et al. 2013; Youssef et al. 2015). However, albeit with metabolic deficiencies, they have been found to be ubiquitous in nature, and mostly distributed in oxygen-limited environments (Castelle et al. 2018; Castelle et al. 2015; Rinke et al. 2013). These studies have revealed their unique and unneglectable roles in biogeochemical cycles (Baker et al. 2020), as well as mysterious interactions with their potential hosts.

*Candidatus* Micrarchaeota and *Candidatus* Parvarchaeota (reassigned to the order *Candidatus* Parvarchaeales within the phylum *Nanoarchaeota* in GTDB r207 [https://gtdb.ecogenomic.org]) were first found in AMD name Archaeal Richmond Mine Acidophilic Nanoorganisms (ARMAN) proposed due to their ultra-small cell and genome sizes (Baker et al. 2010; Baker et al. 2006).

Phylogenetic inference in a later study has shown that *Ca*. Parvarchaeota differs from *Ca*. Micrarchaeota and represent a separate lineage (Chen et al. 2018). Similar to Micrarchaeota, the metabolic potential of Parvarchaeota is versatile with the possession of a diversity of capacities, such as the glycolysis, the tricarboxylic acid (TCA) cycle, the electron transfer chain (ETC), and numerous polysaccharide degradation pathways, which were relatively less seen in other DPANN archaea (Castelle et al. 2018; Chen et al. 2018). The presence of ETC and the TCA cycle in oxygen-limited environments is possibly associated with the consumption of oxygen rather than aerobic respiration. Although initially discovered in AMD, another ARMAN archaea - Micrarchaeota have been tremendously expanded in phylogenetic diversity and have been found to exist in a variety of environments, including extreme habitats such as hot springs and radioactive sites, as well as non-extreme habitats like underground water (He et al. 2021; Sakai et al. 2022; Vázquez-Campos et al. 2021). In contrast, Parvarchaeota appear to inhabit a narrower range of habitats. Thus far, their genomes have been obtained exclusively from acidic and/or thermal environments (Chen et al. 2018; Luo et al. 2020). However, it is still unclear regarding their adaptation mechanisms to harsh conditions and how they evolved to give rise to such metabolic versatility.

Many archaeal lineages were considered to originate from high-temperature (thermal) environments. It’s well supported that the early environment of the Earth was found to be of high temperature and pressure, where only hyperthermophiles (often seen in deep-rooted archaea and bacteria) would have survived such circumstances (Stetter 2006). The growth conditions characterization of microorganisms is essential to understanding their physiology. However, basic growth conditions such as optimal growth temperature (OGT) and optimal pH are inaccessible for archaea and bacteria without cultured representatives. Pieces of evidence also showed optimal growth temperatures (OGTs) of bacteria and archaea can be well explained by the G + C content of rRNAs and amino acid compositions (Galtier and Lobry 1997; Zeldovich et al. 2007; Zhaxybayeva et al. 2009). Consequently, efforts to use genomic composition to reflect environmental conditions have been proposed as “reverse ecology” approaches (Li et al. 2008). In this sense, characteristics such as OGT and pH have been estimated by various methods. For example, the over-representation of seven amino acids Ile, Val, Tyr, Trp, Arg, Glu, and Leu (IVYWREL) were shown to be positively correlated with the OGT of Archaea and Bacteria, and subsequently could be used for OGT prediction (Zeldovich et al. 2007). Other methods were based on amino acid compositions, or a combination of 16S rRNA genes, tRNA and ORFs, and protein sequences (Li et al. 2019; Sauer and Wang 2019). Further ancestral state reconstructions have revealed that many archaeal ancestors adapt to high-temperature environments with high OGTs (Groussin et al. 2013; Groussin and Gouy 2011; Williams et al. 2017).

As one of the major deep-rooted archaeal lineages, the DPANN superphylum was shown to have a mesophilic origin in the previous study (Williams et al. 2017). However, this could be a result of the lacking representation of genomes in high-temperature environments. Thus, it is still unclear whether DPANN as a whole or at least some specific lineages like Parvarchaeota originated from high-temperature environments. In addition, if Parvarchaeota have a distribution uniquely in acidic habitats, whether it was a legacy of early colonization in acidic hot springs or a later adaptation after the dispersal to acidic habitats remains unknown.

Here, we reconstructed 28 metagenome-assembled genomes (MAGs) from hot spring samples of Tengchong, Yunnan, China. With genome-resolved metagenomics analyses and applying comprehensive phylogenetic analyses, we identified 10 MAGs branched deeply within Parvarchaeales, 4 MAGs belong to Parvarchaeum, and 14 MAGs form a new clade that is sister to Parvarchaeales representing a new order according to the GTDB-based taxonomic assignments. We also reconstructed the metabolic potential and conducted in-depth comparative and evolutionary genomics analyses on these genomes, demonstrating their metabolic versatility and adaptation processes to these extreme environments. With the reconstruction of several ancestral genomic features, we sought to examine the possible evolutionary trajectory regarding the adaptation of these lineages to different extreme environments.

## Results

### Genomes quality and taxonomy hierarchy of the newly discovered genomes

A total of 28 metagenome-assembled genomes (MAGs) were reconstructed from 28 metagenomes of geothermal springs sediment samples collected from Jan 2017 to Aug 2020 in Tengchong, Yunnan, China (Supplementary Tables 1 and 2). Three approaches were applied to evaluate the genome quality: 1) the occurrence frequency of genes among 48 single copy genes (SCGs, see Supplementary Table 3 for gene counts of each SCG) (Castelle et al. 2015; He et al. 2021); 2) using the latest published CheckM2 package to assess genome quality (Chklovski et al. 2022); 3) estimating the genome completeness and contamination using CheckM with the exclusion of markers that were universally missing in the corresponding genus (Supplementary Table 4) (Dombrowski et al. 2020). All MAGs were with low contamination (<5%). Twenty MAGs recovered in this study were of high quality (completeness≥ 90%, with the presence of both 16S and 23S rRNA and >18 tRNA) supported by at least one approach. The remaining MAGs were of medium quality with genome completeness > 50%. The results of taxonomy with GTDB-tk (Chaumeil et al. 2019) demonstrated that 14 genomes were affiliated with previously undescribed archaeal order “DTBS01” (herein named *Candidatus* Jingweiarchaeales). Among them, seven MAGs were classified into a potential new genus, while the other seven MAGs were assigned to the undescribed genus “DTBS01”, with the names *Candidatus* Jingweiarchaeum and *Candidatus* Haiyanarchaeum proposed (Supplementary Table 1, Taxonomy appendices).

### Phylogenetic placement

Consistent with the previous study (Dombrowski et al. 2020), the phylogenetic inference based on the concatenation of 53 conserved archaeal marker genes demonstrated that the DPANN superphylum was grouped into two clusters – DPANN cluster I and DPANN cluster II, with high bootstrap confidences (≥ 90, Fig. 1, see counts of all 53 markers in each genome in Supplementary Table 5). During the phylogenetic construction, Huberarchaeota genomes were excluded due to the possible long-branching attraction effect. When all MAGs from Parvarchaeales were considered, the phylogeny placed Huberarchaeota as the sister lineage of Nanoarchaeota (Figure S1a). We further tested if this was possibly due to the uneven selection of representative genomes of these lineages (Venditti et al. 2006). However, when those closely related MAGs were dereplicated using dRep (Olm et al. 2017) with 99% ANI as the cutoff, the phylogenetic position of Nanoarchaeota remains unstable after the selection of dRep (Figure S1b). Specifically, Huberarchaeota branched deeply within the Nanoarchaeota with low bootstrap confidence at their parent node. We further sought to examine if this is caused by the potential long-branch attraction (LBA) effect by the Huberarchaeota genomes since they presented a deep-rooted long branch in previous studies (Bergsten 2005; Dombrowski et al. 2020). After the exclusion of Huberarchaeota genomes, the topology structure became stable and each node within Nanoarchaeota is well supported with bootstrap confidence ≥ 80 (Fig. 1). Two reference genomes that were previously labeled as Parvarchaeota in NCBI taxonomy and order CSSed11-243R1 in GTDB, was shown to form a sister lineage with Jingweiarchaeales. Within the phylum Nanoarchaeota, Nanoarchaeales represent the deepest-rooted lineage, while Jingweiarchaeales, CSSed11-243R1, and Parvarchaeales together form a sister lineage with Woesearchaeales and Pacearchaeales.

**Figure 1.**
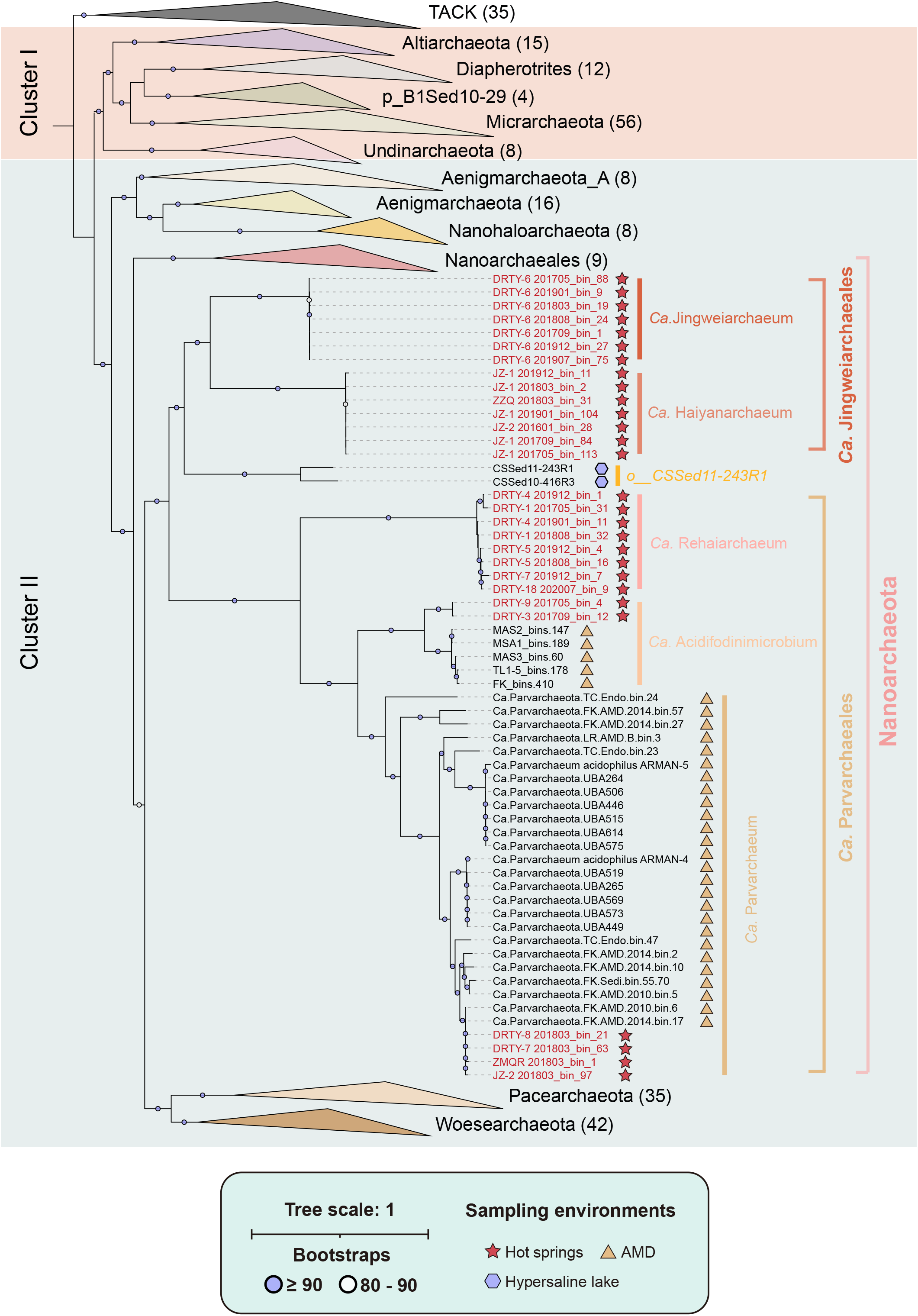
Phylogenetic placement of the newly discovered MAGs. Genomes recovered in this study were labeled in red on the tips of the phylogenetic tree. Sampling environments of the associated genomes were shown in red stars for hot springs, purple hexagons for hypersaline lakes, and yellow triangles for acid mine drainage (AMD) respectively. For bootstraps in each ancestral node, values ≥ 90 were shown in purple solid circles, and values between 80 to 90 were shown in hollow circles.

### Environmental distribution

We found that both Jingweiarchaeales and Parvarchaeales present low relative abundances in the associated sampling environments (<0.5% for Jingweiarchaeales and <4% for Parvarchaeales, Figure S2). A previous pioneering study revealed the limited distribution of Parvarchaeales, which were detected only in acidic environments such as AMD and Tengchong acidic endolithic community (Chen et al. 2018). This led to the question that whether Parvarchaeales is an acidophilic lineage that specifically inhabits acidic environments. In this study, we found that most of the available Parvarchaeum genomes were retrieved from extremely acidic environments with pH <3 and thus might represent an acidophilic archaeal lineage (Figure S2, Supplementary Table 1) (Baker-Austin and Dopson 2007). This interference was consolidated by observation in our samples that Parvarchaeales often appeared with higher relative abundance (>1%) when pH is lower than 4 and increased over time in the site DRTY-6 as pH decreased (Figure S2).

### Metabolic potentials

By reconstructing the metabolic pathways of Jingweiarchaeales and Parvarchaeales (Fig. 2, Supplementary Table 6), Jingweiarchaeum is the only genus with the detection of a complete AMP that was commonly observed in DPANN lineages. Phylogenetic inference identified type III *rbcL* gene encoding the large subunit of ribulose 1,5-bisphosphate carboxylase/oxygenase (RuBisCO) in Jingweiarchaeum, consolidating the potential function involved in nucleotide salvage pathway (Figure S3) (Jaffe et al. 2019; Sato et al. 2007; Wrighton et al. 2016). The detection of a partial pentose phosphate pathway (PPP) in Jingweiarchaeum suggested that they could convert F6P and GAP to PRPP. Given the absence of glucokinase and fructose-1,6-biphosphate kinase (F1,6BP) in glycolysis, the partial PPP coupling with AMP enables the conversion of G3P, which further leads to the generation of pyruvate and ATP by linking to the glycolysis. The TCA cycle and electron transfer chain (ETC), such as the V-type ATPase complex, are beyond detection, suggesting that Jingweiarchaeum may rely heavily on fermentation for energy production. As expected, Jingweiarchaeum might be able to ferment acetate by harboring acetyl-CoA synthetase (ADP-forming) and acetyl-CoA synthetase encoded by *acdAB* and ACSS1_2 (Glasemacher et al. 1997). In contrast, Haiyanarchaeum possess an even more reduced glycolytic pathway and lack PPP and AMP. In addition, genes associated with acetate fermentation are also missing, suggesting a minimal metabolic capacity that is similar to the genome of *Candidatus* Woesearchaeota archaeon AR20, Haiyanarchaeum may be obligate symbionts and heavily rely on their hosts for resources (Castelle et al. 2018; Castelle et al. 2015).

**Figure 2.**
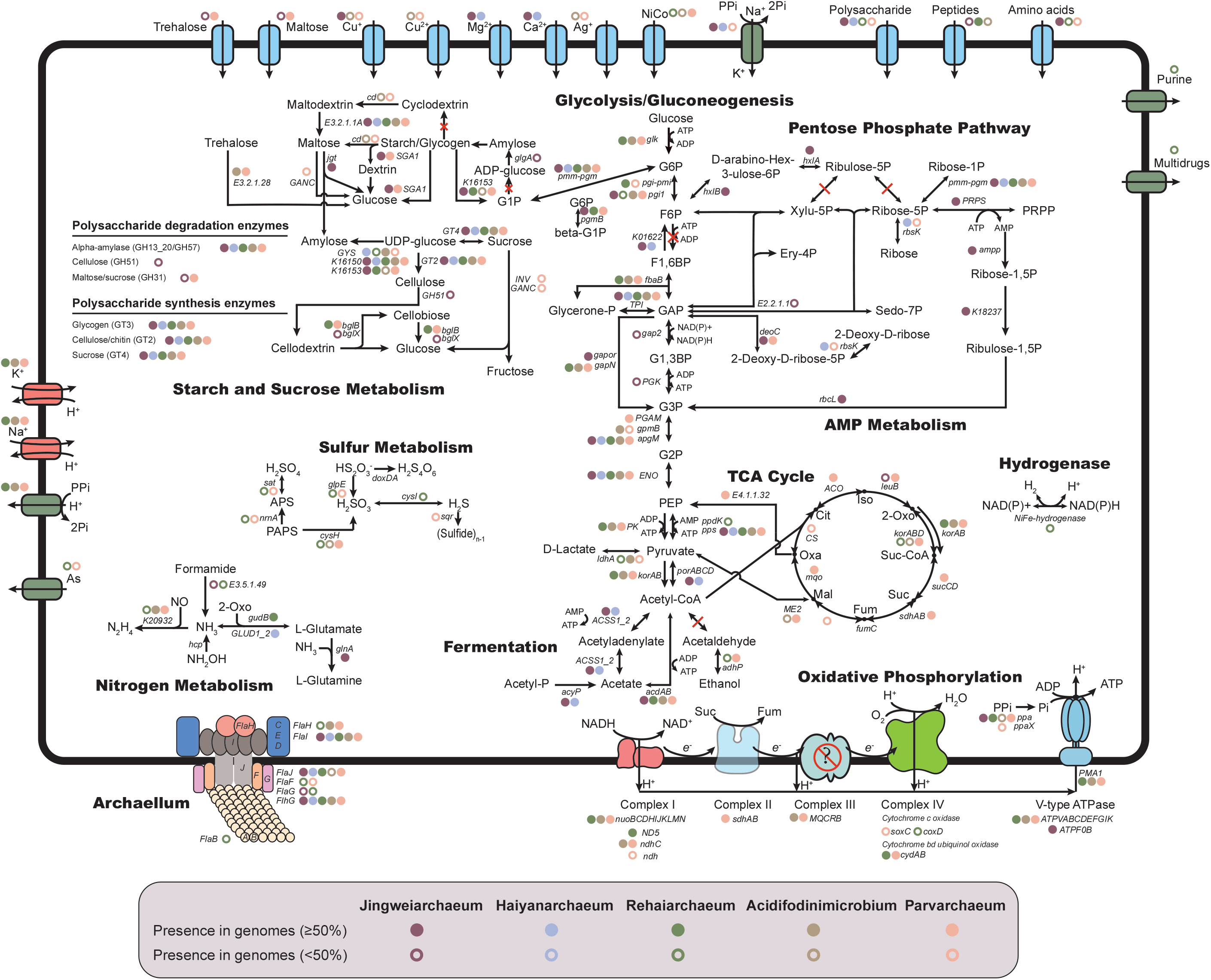
Metabolic pathway in Jingweiarchaeum, Haiyanarchaeum, Rehaiarchaeum, Acidifodinimicrobium, and Parvarchaeum that recovered from 60 MAGs. The presence of genes in Jingweiarchaeum, Haiyanarchaeum, Rehaiarchaeum, Acidifodinimicrobium, and Parvarchaeum were shown in dark-red, blue, green, yellow, and pink symbols, respectively. Genes that are present in more than 50% of the genomes were illustrated in solid circles, while genes present in less than 50% but at least one MAG were shown in hollow circles. The names of genes were labeled next to the circles representing the presence of genes. Abbreviations: G6P, glucose-6-phosphate; F6P, fructose-6-phosphate; F1,6BP, fructose-1,6-bisphosphate; GAP, glyceraldehyde-3-phosphate; Glycerone-P, glycerone-phosphate; G1,3BP, glycerate-1,3-bisphosphate; G3P, glycerate-3-phosphate; G2P, glycerate-2-phosphate; PEP, phosphoenolpyruvate; Xylu-5P, xylulose-5-phosphate; Ribulose-5P, ribulose-5-phosphate; Ribose-5P, ribose-5-phosphate; Sedo-7P, sedoheptulose-7-phosphate; Ery-4P, erythrose-4-phosphate; Ribose-1P, ribose-1-phosphate; PRPP, phosphoribosyl diphosphate; Ribose-1,5P, ribose-1,5-bisphosphate; Ribulose-1,5P, ribulose-1,5-bisphosphate; 2-Deoxy-D-ribose-5P, 2-Deoxy-D-ribose 5-phosphate; 2-Deoxy-D-ribose, 2-deoxy-D-erythro-pentose; D-arabino-Hex-3-ulose-6P, D-arabino-3-Hexulose 6-phosphate; G1P, glucose-1-phosphatase; ADP-glucose, adenosine diphosphoglucose; UDP-glucose, uridine diphosphate glucose; Oxa, oxaloacetate; Cit, citrate; Iso, isocitrate; 2-Oxo, 2-oxoglutarate; Suc-CoA, succinyl-CoA; Suc, succinate; Fum, fumarate; Mal, malate; APS, Adenosine 5’-phosphosulfate; PAPS, 3’-Phosphoadenosine 5’-phosphosulfate; TCA cycle, tricarboxylic acid cycle.

Consistent with the previous study, Parvarchaeum (former *Ca*. Parvarchaeota) MAGs encode the near-complete pathways for glycolysis, TCA cycle, and ETC (Chen et al. 2018). However, we were unable to detect phosphofructokinase in all Parvarchaeum MAGs including genomes described in the previous research (Chen et al. 2018). It is unclear whether this is due to annotations with different databases or the changes in associated databases that have altered the previous annotation results. On the contrary, most genes associated with the TCA cycle and ETC were missing in Rehaiarchaeum and Acidifodinimicrobium, indicating that they might rely on acetate fermentation for energy production with the presence of *acdAB* gene (Reeves et al. 1977). Besides, Rehaiarchaeum and Parvarchaeum possess genes encoding alcohol dehydrogenases (*adhP*), enabling them to ferment ethanol. However, none of these MAGs have aldehyde dehydrogenase for the conversion of acetyl-CoA to aldehyde, indicating they might rely on their hosts to promote the fermentation process.

The presence of only one menaquinol-cytochrome c reductase cytochrome b subunit (MQCRB) in Acidifodinimicrobium and Parvarchaeum genomes raise the question of whether these genomes indeed encode fully functional complex III of ETC. Similarly, only two subunits *coxD* and *soxC* involved in complex IV were detected in a few Parvarchaeum MAGs. However, both Rehaiarchaeum and Parvarchaeum have *cydAB* gene encoding cytochrome bd ubiquinol oxidase, which was reported to have high oxygen affinity and was expressed under oxygen-limited conditions (Borisov 2011). The cytochrome bd ubiquinol oxidase could use ubiquinol as an electron carrier coupled with the reduction of oxygen and generate proton motive force (PMF), possible in place of the function of complex III & IV and consume extra oxygen to maintain low-oxygen intracellular environments (Poole and Hill 1997; Tyson et al. 2004). All these might suggest that although with a complete TCA cycle and potentially complete ETC, Parvarchaeum may be oxygen-tolerant that inhabit environments with lower levels of oxygen rather than favoring aerobic environments. However, we could not rule out the possibility that some DPANN may acquire the ability to respire oxygen using the ETC coupling with TCA. Or at least they may use the PMF of their potential hosts when in proximity to the presence of complex V (Castelle et al. 2018).

### Gluconeogenesis, Polysaccharides degradation, and biosynthesis

Although Parvarchaeales lack a complete glycolysis pathway, Jingweiarchaeales MAGs appear to encode genes for the gluconeogenesis pathway with the G6P as the end-product., including *porABCD, pps*, and K01622. The ancestral gluconeogenic enzyme fructose 1,6-bisphosphate (FBP) aldolase/phosphatase exhibiting both the FBP aldolase and FBP phosphatase activity was detected. It’s reported that the FBP phosphatase is irreversible which renders gluconeogenesis unidirectionally (Say and Fuchs 2010). The production of G6P via gluconeogenesis might further be connected to the synthesis of polysaccharides by further conversion of G6P to G1P by *pmm-pgm* gene encoded phosphomannomutase/phosphoglucomutase. Functional annotation results based on comparison to the Carbohydrate-active enzymes (CAZy) database (Lombard et al. 2014) revealed that both Jingweiarchaeales and Parvarchaeales are capable of synthesizing glycogen (β-glucosidase, GT3), cellulose (β-glucosidase, GT2), and sucrose (sucrose synthase, GT4). Besides, all genera contain genes encoding glycogen degradation including GH13_20 and/or GH57. The detection of SGA1 and *jgt* encoding glucoamylase and 4-alpha-glucanotransferase in Jingweiarchaeum suggest that this genus is capable of degrading maltose with the release of glucose. It seems plausible that maltose can serve as an intermediate of starch/glycogen degradation which leads to the final production of monosaccharides. This could be further exemplified by the observation of GH31 in Jingweiarchaeum MAGs. In addition, Jingweiarchaeum can also degrade cellulose with GH51, demonstrating its versatility in polysaccharide utilization. The presence of trehalose and maltose transport systems, as well as genes associated with their degradation via alpha,alpha-trehalase (EC: 3.2.1.28) in Parvarchaeum, suggests that the external import may serve as an alternative source of sugars to sustain the glycolysis pathway in this genus. Taken together, we conjectured that the detected gluconeogenesis and polysaccharide biosynthesis pathways in Jingweiarchaeales. Especially Jingweiarchaeum might contribute to the storage of polysaccharides under nutrient-rich conditions. Both of these DPANN archaea and putative hosts may gain ATP from these monosaccharides to sustain their growth through the glycolytic pathway or the aforementioned coupled pathway of partial glycolysis, AMP and PPP.

### Cell membrane biosynthesis

All three genera in Parvarchaeales lack genes to biosynthesize isoprenoid (IPP) and phospholipid backbone, the main constituents of the cell membrane. Similarly, the related genes also failed to be detected in most Jingweiarchaeales MAGs except *araM* gene, which encodes glycerol-1-phosphate dehydrogenase [NAD(P)^+^] (G1PDH). To our surprise, each Jingweiarchaeum MAG possesses a *mvk* gene that encodes mevalonate kinase, which is vital to the biosynthesis of archaeal membrane precursor IPP. However, the biosynthetic pathway of IPP and further to archaeol was incomplete in Jingweiarchaeum due to the lack of genes encoding phosphomevalonate kinase (E2.7.4.2), diphosphomevalonate decarboxylase (*mvd*), and archaetidylserine synthase (*pssA*). Haiyanarchaeum on the other hand have even more genes missing in the associated pathways. Consequently, they must rely on their hosts to provide key intermediates to assist the biosynthesis of archaeal cell membranes. Although the *glpA* gene encoding glycerol-3-phosphate dehydrogenase (G3PDH) was surprisingly found in all genera of Parvarchaeales, they are unable to synthesize bacterial cell membrane since most genes associated with the MEP pathway and phospholipid biosynthesis were absent. All these results indicated the host dependence of Jingweiarchaeales and Parvarchaeales.

Archaellum and pilus are considered to be associated with the motility, attachment, and biofilm formation of archaea (van Wolferen et al. 2018). Several previous studies have demonstrated that DPANN archaea probably can penetrate their host cells via pili or pili-like structures (Comolli and Banfield 2014; He et al. 2021). We found that besides *flaHIJ* genes encoded for archaella base structure, the *flaB* gene encoding major archallellin and other genes were found in one Rehaiarchaeum MAG DRTY-4_201901_bins_11. Besides, genes encoding pilin (*pilA*-subfamily, arCOG02420 for Jingweiarchaeum and arCOG03871 for Haiyanarchaeum) were detected in all genomes, as well as genes encoding other structural proteins in some of the MAGs within Jingweiarchaeales and Parvarchaeales (Makarova 2016).

### Hydrogen, Sulfur and Nitrogen metabolism

One Rehaiarchaeum MAG, DRTY-1_201705_bins_31, possesses NiFe-hydrogenase encoded by *hyaAB*, as well as several hydrogenase maturation and formation proteins. The absence of hydrogenase indicates that most members of Jingweiarchaeales and Parvarchaeales rely on alternative strategies for NAD(P)H and NAD(P)^+^ interconversion. Unlike most DPANN archaea, Parvarchaeales genomes harbor several genes related to sulfur and nitrogen metabolism.

Specifically, Rehaiarchaeum harbor the *cysI* gene encoding sulfite reductase (NADPH) (Zeghouf et al. 2000). The reversible reaction may result in the interconversion of NAD(P)+ and NAD(P)H in place of the limited hydrogenases, which may couple with their organic carbon oxidation in both partial glycolysis and acetate/ethanol fermentation. Strikingly, genes encoded for hydrazine synthase (K20932, EC 1.7.2.7) were found in all three genera within Parvarchaeales which has never been found in any DPANN archaea. As reported, this enzyme participates in anaerobic ammonium oxidation by the production of the highly reactive hydrazine from ammonia and nitric oxide, which could be further reduced to N_2_ (Dietl et al. 2015; Kartal et al. 2011). However, the exact role of this gene in Parvarchaeales still needs to be experimentally verified.

### Comparative genomics analysis on the adaptabilities to various habitats

To ensure the accuracy of genome statistics and comparative genomics, only genomes in this study and reference genomes with ≥ 80% completeness determined both by 48 SCGs and CheckM2 were kept for further analyses. However, with the assessment of genome quality by CheckM2 and occurrence frequency of markers among 48 SCGs, we observed that the Rehaiarchaeum and Acidifodinimicrobium MAGs have collectively low genome qualities. Thus, we further examine if there were other markers universally missing in these lineages. Apart from PF01849.13, PF01912.13, PF01922.12, PF04127.10, PF05221.12, PF06026.9, TIGR00336, TIGR00670, and TIGR01213 that were consistently absent in Jingweiarchaeales and Parvarchaeales, several other markers were also missing within each specific genus. For example, the three genera in Parvarchaeales lack PF13685.1, TIGR03677, and TIGR00432, while they all can be detected in Jingweiarchaeales. We further modified the 149 archaeal markers used by CheckM1 by excluding markers that were universally missing in each genus, which include 11 markers missing in Jingweiarchaeum, 21 markers in Haiyanarchaeum, 29 markers in Rehaiarchaeum, 21 markers in Acidifodinimicrobium, and 17 markers in Parvarchaeum to assess genome quality of each MAG (Supplementary Table 5). By applying this approach, 48 MAGs with completeness ≥ 80% were kept for further comparative genomics analysis (Supplementary Table 1). Due to the lack of complete genomes, however, we still cannot rule out the possibility that technical biases during the sequence assembly and genome binning may lead to missing markers.

To avoid bias of genomic feature statistics caused by incomplete genome assembly, we examined the effect of genomic completeness on the relative genome size differences in all genera in this study, which showed to be stable over different approaches (Fig. 3a, Figures S4a-c). Overall, Jingweiarchaeum MAGs have the largest genome sizes among all five genera and followed by Parvarchaeum, while Rehaiarchaeum have the smallest genome sizes. This is significantly better explained by the number of protein-coding sequences (CDSs), where a significant positive relationship was observed between genome sizes and the number of CDSs (adj. R^2^ = 0.94, p = 3.1324e-31, Figs. 3b and 3g). Genomes of Haiyanarchaeum appear to be the most compact, with small genome sizes, high coding density, and the highest ratio of overlapping genes (Figures S4e and 4f). The high compactness of these genomes might be a result of adaptation to environments with high temperatures (Fig. 3c) (Sabath et al. 2013).

**Figure 3.**
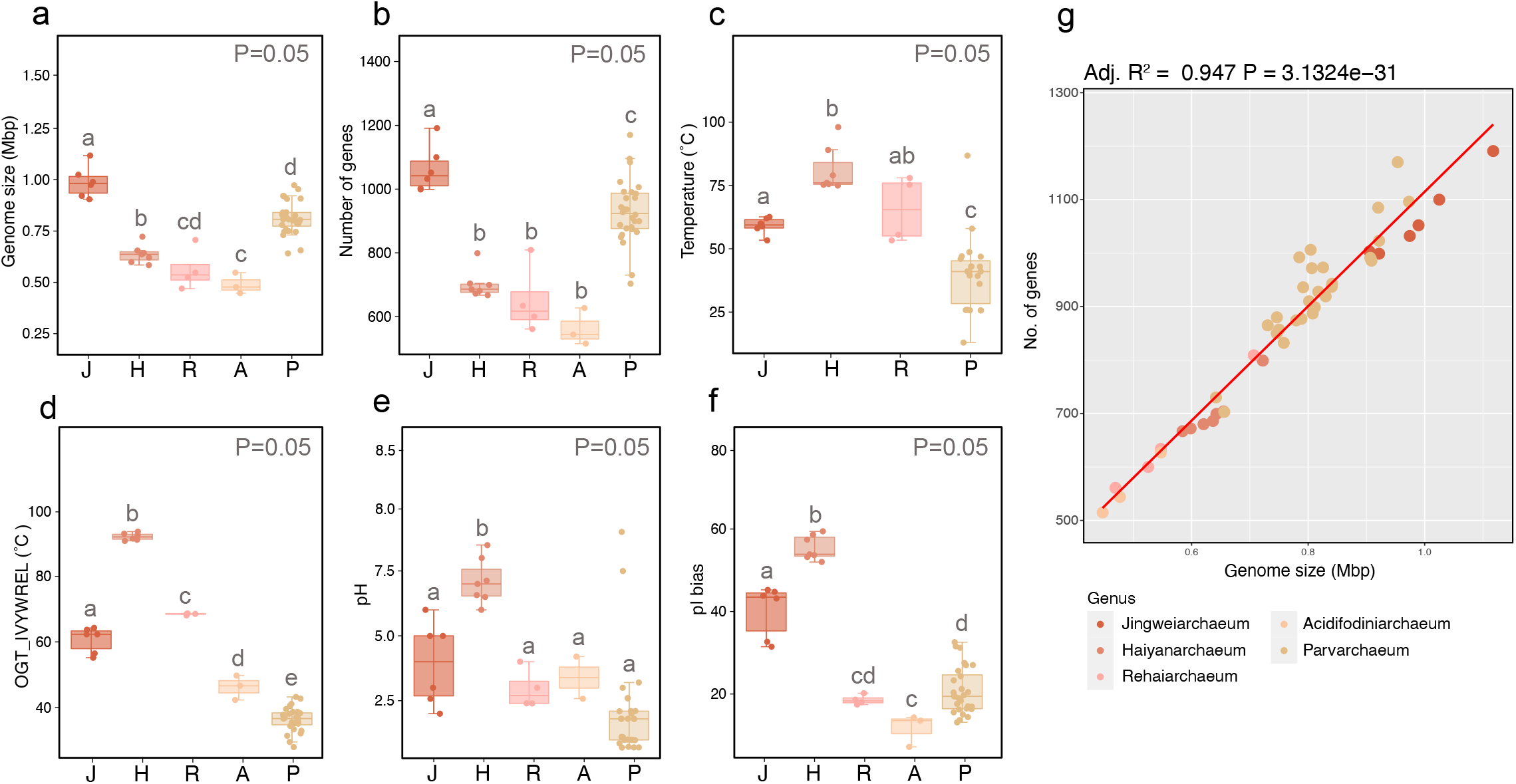
Genomic comparisons of Jingweiarchaeum, Haiyanarchaeum, Rehaiarchaeum, Acidifodinimicrobium, and Parvarchaeum. Differences of **a)** genome sizes, **b)** the number of genes, **c)** temperature, **d)** OGT predicted by the overrepresentation of seven amino acids (IVYWREL), **e)** pH, and **f)** average isoelectric point (*pI*), and were visualized using ggplot2 package (v3.3.6) (Wickham 2016b). Different alphabets show significant differences between groups with P <0.005.

Regarding the GC content of genomes of these genera, they vary across lineages (Figure S4g). Principle coordinate analyses (PCoA) based on the metabolic profiles of KEGG and arCOG annotation results demonstrated that genomes were well clustered based on their phylogenetic position rather than habitats (Figure S5). The close distance between Acidifodinimicrobium and Parvarchaeum suggests that a recent geographical dispersion has possibly occurred. This is also supported by the average amino acid identities between these lineages (Figure S6). The larger sizes of Parvarchaeum within the PCoA plots exhibit higher genomic plasticity among these genera, which may assign them the ability to better adapt to different habitats. However, the clustering of Parvarchaeum genomes from both high and low temperature environments suggests that the adaptation to different environments may have not yet induced notable genomic shifts in time. This inference can be further corroborated by the observation of a wider range of GC content, isoelectric point (*pI*), and estimated optimal growth temperatures (OGTs) in this genus, which demonstrates a more diverse nucleotide/amino acid composition (Figs. 3d and 3f, Figures S4g-j). In contrast, Jingweiarchaeum, Haiyanarchaeum, and Rehaiarchaeum primarily inhabit thermal habitats, whereas Acidifodinimicrobium and Parvarchaeum favor environments with lower temperatures.

### Adaptation to high-temperature environments

The investigation of sampling sites suggested that Haiyanarchaeum were mostly derived from samples with higher temperatures, while Parvarchaeum were from samples with lower temperatures (Fig. 3c). Overall, Haiyanarchaeum appear to be hyperthermophilic with the OGTs > 80 °C predicted by IVYWREL (Fig. 3d). Expectedly, *rgy* genes encoding reverse gyrase that was ubiquitously presented in hyperthermophiles were detected in genomes of Haiyanarchaeum, further suggesting that they might indeed represent a hyperthermophilic lineage (Supplementary Table 6) (Heine and Chandra 2009; Kampmann and Stock 2004; Lulchev and Klostermeier 2014). Given the exclusive detection of currently available Jingweiarchaeum MAGs we infer that Jingweiarchaeum might be also thermophiles. Most MAGs within Parvarchaeum and Acidifodinimicrobium are mesophiles, of which growth temperatures below 50 °C are preferable.

However, when looking into other genes previously considered vital to the thermal adaptation (Lewin et al. 2013), they didn’t show specificity to Jingweiarchaeales that of higher OGTs and sampling temperatures (Supplementary Table 6). These genes encoding heat shock proteins (HSPs), chaperones, and prefoldins which can protect proteins from denaturation were widely detected among all MAGs (Fukui et al. 2005). Besides, the possession of DNA repair protein RadA that mediates the recombination process (Beam et al. 2002), topoisomerases that maintain DNA structure, and polyamines such as spermidines (*speE*) that stabilize the cytoplasm also confer the ability to resist high-temperature stress (Fukui et al. 2005). Save for thermal adaptation, these genes can also allow them to survive under other stresses, such as acid stress.

The equivalent distribution of these genes among Jingweiarchaeales and Parvarchaeales MAGs strengthens this inference. It is noteworthy that Jingweiarchaeales genomes uniquely harbor DNA repair systems encoded by ERCC4, *ykoV*, and *splB*. However, we could not confirm whether they were exploited to cope with high-temperature stress.

The predicted OGT in this study largely reflects the preference of amino acid usage for each genome, since all methods consider only or partly the amino acids compositions of proteomes for prediction. As expected, the accumulation of glutamate (Glu, E) was observed in the genomes of Jingweiarchaeum, Haiyanarchaeum, and Rehaiarchaeum (Figure S7). Haiyanarchaeum in particular, also with more representation of arginine (Arg, R). Previously studies have shown that the higher fraction of Arg residues could enhance thermostability with more stable salt bridges (Siddiqui et al. 2006), among which the combination between arginine and glutamate (R+E) was shown to be the strongest (White et al. 2013). The accumulation of proline could also potentially lower the flexibility of protein secondary structure as well as lower the efficiency of proline isomerization, which may also increase the thermostability of proteins (Feller 2013; Najmanovich et al. 2000; Sakaguchi et al. 2007). As a result, the higher fractions of R+E and P within Haiyanarchaeum proteomes may be vital to their adaptation to higher temperatures environments. Collectively, the adaptation strategies of Jingweiarchaeales, especially hyperthermophilic Haiyanarchaeum, may include genome streamlining with small genome sizes, high coding density, high ratio of overlapping genes, and bias of amino acid usages that favor high-temperature environments.

### Acidic adaptation

Many MAGs from Parvarchaeales in this study were recovered from acidic environments, such as AMD and acid hot springs. Given this, we further examine the *pI* of each genome to reveal the intracellular pH (Kozlowski 2017; Kozlowski 2016). Interestingly, all genomes, regardless of the pH of belonging samples exhibit bimodal distributions with a relatively low fraction of proteins with *pI* close to neutral pH (Figures S8-13). This is expected to be a result of the instability and insolubility of proteins in pH close to *pI* (Kiraga et al. 2007). Further, as shown by the *pI* bias of these genomes, none of these genomes were over-represented by acidic proteins than basic proteins (*pI* bias > 0) demonstrating that the inner cell environments of these archaea were near-neutral (Fig. 3f, Figure S4h). Unlike most halophilic archaea which sustain low acidic intracellular pH to improve the amino acid solubility (Kiraga et al. 2007), the maintenance of near-neutral intracellular pH suggested that Parvarchaeales tend to be acid-tolerant microbes rather than acidophiles. This inference is further supported by the detection of potassium/hydrogen antiporter (*cvrA*), P-type H^+^-exporting transporter (PMA1), and Na+/H+ antiporter in Parvarchaeales genomes, which could potentially export excess protons extracellularly to alleviate the acid stress (Fig. 2, Supplementary Table 6). Interestingly, Jingweiarchaeum appear to have different strategies to cope with acidic stress including the decarboxylase of organic acids (*speA, mfnA, pdaD*) and the formation of a proton-impermeable membrane by the synthesis of hopanoid (*sqhC*) to maintain the intracellular pH homeostasis (Baker-Austin and Dopson 2007).

Oxidative stress appears to be another obstacle to survival in acidic environments (Ram et al. 2005). As aforementioned, most MAGs within Parvarchaeales harbor the high oxygen affinity cytochrome oxidase *cydAB*, which might be associated with the removal of oxygen (Borisov 2011; Poole and Hill 1997; Tyson et al. 2004). Besides, other genes including superoxide dismutase (*sodA*), and peroxiredoxin (*bcp* and *ahpC*) were solely found in all genera in Parvarchaeales. These genes guarantee the physiological activity of these microbes when they are exposed to oxic environments.

### Origin and evolution of Jingweiarchaeales and Parvarchaeales lineages

By applying three approaches to calculate the OGTs of all MAGs in this study, we observed that three deeply rooted genera including Jingweiarchaeum, Haiyanarchaeum, and Rehaiarchaeum are more likely to inhabit thermal environments (Figs. 3c and 3d, Figures S4i and 4j). With the deep-rooted lineages of Jingweiarchaeales and Parvarchaeales showing high OGTs, we hypothesized that these lineages possibly originated from high-temperature environments. This inference is well supported by the OGT prediction of ancestral nodes. OGTs predicted by IVYWREL reveal that the last common ancestor of Jingweiarchaeales and Parvarchaeales (LJPCA) have an OGT of 66.09°C (Fig. 4a). After the differentiation of LJPCA, OGTs tend to decrease toward the evolution of Parvarchaeum, but was with an increasing pattern in Jingweiarchaeum. The last common ancestors of Parvarchaeum (LPaCA) and Haiyanarchaeum (LHaCA) may thrive with OGTs of 37.67°C and 93.24°C respectively. In contrast, OGTs of the last common ancestors of Jingweiarchaeum and Rehaiarchaeum (LJiCA and LReCA) remained relatively unchanged compared to LJPCA. Collectively, both phylogenetic placements and OGT prediction of ancestral nodes support the hot origin of two orders. Further evolution resulted in the adaptation of Parvarchaeum to the mesophilic environments and the emergence of hyperthermophilic Haiyanarchaeum.

**Figure 4.**
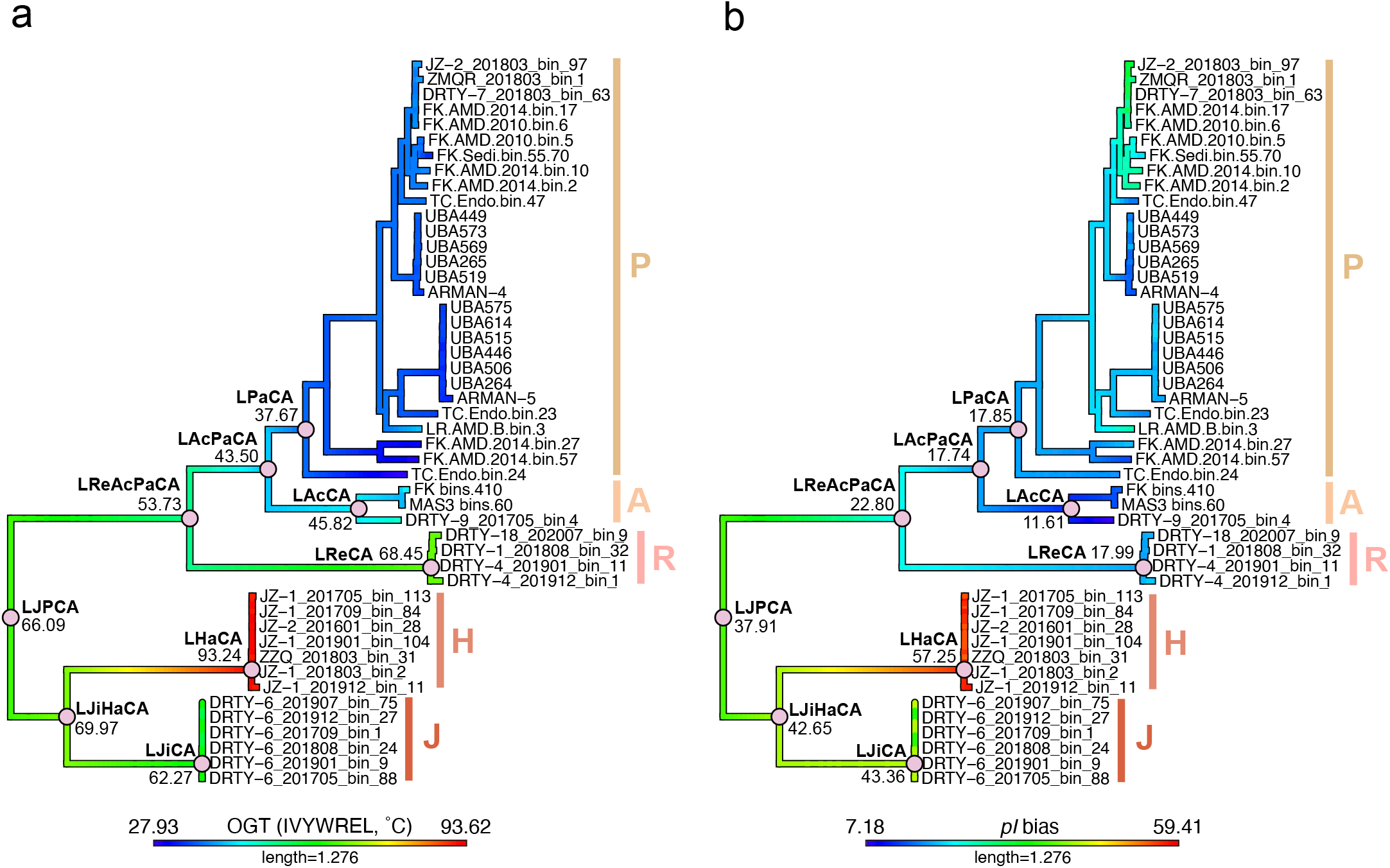
Inference of ancestral traits of Jingweiarchaeales and Parvarchaeales including predicted optimal growth temperature (OGT) and isoelectric point (p*I*). The evolutionary changes of a) OGT predicted by IVYWREL, b) *pI* bias from the common ancestor of Jingweiarchaeales and Parvarchaeales (LJPCA) were shown along with the topology of the phylogeny placement in Fig 1. The nodes of the inferred last common ancestors were marked by pink solid circles, with the predicted genomic feature of the last common ancestor labeled below. Abbreviations: LJPCA, the last common ancestor of Jingweiarchaeales and Parvarchaeales; LJiHaCA, the last common ancestor of Jingweiarchaeum and Haiyanarchaeum; LReAcPaCA, the last common ancestor of Parvarchaeum, Acidifodinimicrobium and Rehaiarchaeum; LJiCA, the last common ancestor of Jingweiarchaeum; LHaCA, the last common ancestor of Haiyanarchaeum; LAcPaCA, the last common ancestor of Acidifodinimicrobium and Parvarchaeum; LPaCA, the last common ancestor of Parvarchaeum; LAcCA, the last common ancestor of Acidifodinimicrobium; LReCA, the last common ancestor of Rehaiarchaeum. The color bar below each figure illustrates the color ranges corresponding to the values of each trait.

To uncover the amino acid usage pattern across the two orders, the ancestral state reconstruction of *pI* and *pI* bias was estimated (Fig. 4b, Figures S4h). Generally, all leaves and ancestral nodes neither show low average *pI* nor low *pI* bias values, suggesting that they sustain a circumneutral intracellular pH. Though all nodes span a narrow range of *pI*, we still observe an increasing pattern from the LJPCA to the LJiCA but the opposite trend from LJPCA to the LPaCA. The variation is more notable with regard to pI bias calculation. To cope with acid stress and manage a circumneutral intracellular pH, different strategies have been adopted. As aforementioned, Parvarchaeales tend to purge protons with proton efflux systems, while Jingweiarchaeales are favored to utilize buffer molecules to maintain intracellular pH homeostasis.

Analyses of the metabolic potentials of Jingweiarchaeales and Parvarchaeales have shown a clear niche differentiation between them. Within Jingweiarchaeales, most microbes are strict anaerobes with fermentation and substrate salvage capacities. Particularly, Jingweiarchaeum harbor incomplete “rerouted glycolysis” that utilized PPP for intermediates conversion and a complete AMP pathway for energy generation. Besides, they are able to obtain energy through acetate fermentation. In contrast, the genomes of Haiyanarchaeum are more streamlined (Fig. 3a and Figures S4e and 4f) with the absence of most genes with regards to glycolysis, AMP, and PPP, suggesting they have to depend on hosts to provide necessary substrates for energy generation. All members in Haiyanarchaeum represent hyperthermophiles due to the wide detection of *rgy* genes. Further investigation of this gene demonstrates that the hyperthermophilic feature in Haiyanarchaeum is acquired from Crenarchaeota via HGT (Figure S14). We reasoned that temperature may lead to the differentiation of these two genera and higher temperature may result in more streamlined genome sizes by losing more non-essential genes.

Parvarchaeales seem to inhabit habitats with relatively low temperatures, though some of the lineages may originate from thermal environments (i.e., Rehaiarchaeum; Fig. 4a). Unlike Jingweiarchaeales which rely on fermentation to gain energy, all genera in Parvarchaeales harbor complete or near-complete glycolysis pathway. Within each order, it seems that the higher temperature the microbes inhabit, the smaller their genomes tend to be. This could be exemplified by Parvarchaeum which optimally grow in the lowest temperature but have the biggest genome size among Parvarchaeales, and the signification positive negative correlation of Jingweiarchaeales MAGs between OGT and genome sizes (Figure S16). Particularly, Parvarchaeum is the only genus that evolved the near complete glycolysis, TCA, and ETC for energy production. In addition, several genes including *sqr, sat*, and *cysH* were detected in Parvarchaeum, suggesting sulfur metabolism may play vital roles during energy generation and substrate cycling. Taken together, we conjecture that oxygen may have driven the genomic diversification within Parvarchaeales.

Finally, notable genomic differences were observed between the two orders. The capacity of fermentation is more prevalent in deep-branching lineages, which were also identified to be anaerobes. Considering that the emergence of life may predate the occurrence of the great oxidation event (GOE), anaerobic respiration is likely to be ancestral across different lineages of life (Knoll and Nowak 2017). Within DPANN archaea, it has also been shown that the ETC pathway may be absent in the common ancestor of DPANN (Beam et al. 2020). Another previous study on Parvarchaeales has indicated the TCA cycle was potentially acquired via horizontal gene transfer (HGT) (Luo et al. 2020). Further phylogenetic inferences revealed that HGT also plays a vital role in the dissemination of genes associated with oxygen tolerance (Figures S17-19). Different origins of *sodA* genes were observed in different members. The *sodA* genes in Acidifodinimicrobium and Parvarchaeum seem to be acquired from Thermoplasmatota, while Rehaiarchaeum likely evolved this gene from Sulfolobales (Figure S17). The phylogeny of *ahpC* gene suggested that Marsarchaeales might be the potential donor (Figure S18). Likewise, many of the thioredoxin-dependent peroxiredoxins encoded by *bcp* in Jingweiarchaeales and Parvarchaeales, are also likely to be transferred horizontally from Marsarchaeales. Few *bcp* genes seem to have bacterial origins (Figure S18). All these demonstrate anaerobic respiration might be ancestral at least for Jingweiarchaeal and Parvarchaeal lineages.

## Conclusion

In this study, we have revealed a new order -Jingweiarchaeales containing two previously undescribed genera Jingweiarchaeum and Haiyanarchaeum that is closely related to Parvarchaeales. With the analyses of metabolic potentials, we revealed that functional differentiation occurred between Jingweiarchaeales and Parvarchaeales. Microbes from Jingweiarchaeales rely mostly on a substrate salvage and fermentative lifestyle to harness the energy, while functional capacities of Parvarchaeales are more versatile with the possession of near complete glycolysis, TCA cycle, and ETC, etc. Comparative genomics demonstrated that Jingweiarchaeales represent a thermophilic lineage, and Haiyanarchaeum in particular may represent a hyperthermophilic genus. Further analyses revealed that the thermal adaptation of these lineages might rely on innate genomic features such as the usage of specific amino acids, genome streamlining, and hyperthermal featured genes such as *rgy*. The adaptation to the acidic environments of Jingweiarchaeum and Parvarchaeales requires the acquisition of genes either associated with proton export, stabilizing of inner cell environments via impermeable cell membrane, or decarboxylation of organic acids. Specifically, the niche expansion of Parvarchaeales was also driven by oxygen stress. By the prediction of OGTs and *pI* and ancestral reconstruction of ancestral features, we demonstrated that both orders have a hot origin, and genomic expansion driven by HGT has led to the adaptation to acidic-mesothermal environments. This study provides insight into the adaption and evolution of DPANN archaea in various extreme environments.

## Methods

### Sampling, DNA extraction, and sequencing

All 28 hot spring sediments samples were collected from Tengchong, Yunnan Province, China in January 2016; May and September 2017; March and August 2018; January, July, and December 2019; and August 2020. DRTY-1, DRTY-3, DRTY-4, DRTY-5, DRTY-6, DRTY-7, DRTY-8, DRTY-9, and DRTY-18 samples were collected from DiReTiYanQu (DRTY), which is an artificial concrete hot spring landscape experiencing area. ZMQR sample was collected from the right side of Zimei spring, a boiling hot spring. ZZQ sample was collected near a boiling hot spring called Zhenzhu spring. These samples were all collected from Rehai National Park, Tengchong, Yunnan, China (24.95 N, 98.44 E). JZ-1 and JZ-2 were sampled from 2 concrete cubic hot spring water pools called JinZe Hot Spring Resort (25.44 N, 98.46 E). The information in detail about all 28 samples, including sampling dates, pH, temperatures and other geochemical properties are available in Supplementary Table 2. We collected the top 1cm of sediment from each site with a sterile iron spoon and transferred these samples to a 50-ml centrifuge tube. All sediment samples were then stored in liquid nitrogen before transporting to the lab. Samples were stored at -20°C in the lab until DNA extraction.

Community DNA was extracted using the PowerSoil DNA isolation kit (MoBio), from about 15 grams of sediment in each sample. We used an M220 Focused-ultrasonicator NEBNext and an Ultra II DNA library prep kit to build libraries with an insert size of 350 bp. The concentration of genomic DNA was measured with a Qubit fluorometer. The total genomic DNA was sequenced with an Illumina Hiseq 4000 instrument at Beijing Novogene Bioinformatics Technology Co., Ltd. (Beijing, China). Each sample was used to generate 30 Giga base pairs (Gbp) of raw sequencing data.

### Metagenomic assembly and genome binning

Raw sequencing data were preprocessed following the previously described workflow (Hua et al. 2018) to remove replicated reads and trim low-quality bases. All quality reads of each dataset were *de novo* assembled by using SPAdes v3.9.0 (Bankevich et al. 2012), with the parameters: -k 21,33,55,77,99,127 –meta. In each assembly, scaffolds with a length of <2,500 bp were removed. BBMap v38.92 (http://sourceforge.net/projects/bbmap/) was used to calculate the coverage information by mapping clean reads to corresponding assembled scaffolds without cross-mapping using the parameters of k=15 minid = 0.97 build = 1. Three software, CONCOCT (v.1.1.0) (Alneberg et al. 2014), Maxbin2 (v.2.2.7) (Wu et al. 2016), and MetaBAT (version 2.12.1) (Kang et al.) were used for automated binning to generate candidate bins for each sample. The best bins were selected by using DAS Tool (v.1.1.3), based on the single-copy genes (SCGs) scoring rank of predicted completeness and contamination of each bin (Sieber et al. 2018). To further improve the quality of genomes, all generated bins were mapped with quality reads of the corresponding metagenome with BBMap. Then mapped reads of each bin were reassembled with SPAdes following the parameters: -k 21,33,55,77,99,127 --careful. CheckM v1.1.3 (Parks et al. 2015) was used to estimate the contaminations and strain heterogeneity of reassembled bins, while the metric completeness was calculated with modified CheckM, which is mentioned next. Moreover, dRep (v3.2.2) (Olm et al. 2017) was used to dereplicate all 61 associated genomes in this study at 99% ANI (strain level), and 21 genomes were selected for phylogenetic reconstruction in Figure S1b. In addition, the relative abundance of each lineage in Figure S2 was calculated based on the coverage of the 21 representative genomes across metagenomic samples with occurrences of lineages of this study.

### Genome quality assessment

Genome qualities of the MAGs were assessed with three methods: 1) 48 previously described single copy genes (SCGs) (He et al. 2021). 2) CheckM2 that includes models for the genome quality assessment of DPANN superphylum archaea (Chklovski et al. 2022). 3) CheckM1 with the exclusion of markers that were missing in all genomes of each genus. For CheckM1, nine of the 149 universal markers of archaea were missing in the genomes of all the lineages in this study including PF01849.13, PF01912.13, PF01922.12, PF04127.10, PF05221.12, PF06026.9, TIGR00336, TIGR00670, and TIGR01213. For each specific genus, nine markers were missing Jingweiarchaeum (PF01849.13, PF01912.13, PF01922.12, PF04127.10, PF05221.12, PF06026.9, TIGR00336, TIGR00670, TIGR01213); 21 were missing in Haiyanarchaeum (PF00398.15, TIGR02076, PF06418.9, PF01725.11, PF04019.7, TIGR00270, TIGR00057, PF00832.15, TIGR00549, PF01982.11, PF00958.17, PF01864.12, PF01849.13, PF01912.13, TIGR00670, PF06026.9, PF01922.12, PF04127.10, PF05221.12, TIGR00336, TIGR01213); 29 were missing in Rehaiarchaeum (PF00900.15, PF00466.15, TIGR02338, PF04010.8, TIGR00422, TIGR00344, PF06418.9, PF01282.14, PF01725.11, PF04019.7, PF00831.18, TIGR00057, TIGR03677, TIGR00432, PF13685.1, PF00832.15, TIGR00549, PF01982.11, PF00958.17, PF01864.12, PF01849.13, PF01912.13, TIGR00670, PF06026.9, PF01922.12, PF04127.10, PF05221.12, TIGR00336, TIGR01213); 24 were missing in Acidifodinimicrobium (PF02005.11, PF08071.7, TIGR02076, PF01725.11, PF04019.7, PF00831.18, TIGR00057, TIGR03677, TIGR00432, PF13685.1, PF00832.15, TIGR00549, PF01982.11, PF00958.17, PF01864.12, PF01849.13, PF01912.13, TIGR00670, PF06026.9, PF01922.12, PF04127.10, PF05221.12, TIGR00336, TIGR01213); and 17 were missing in Parvarchaeum (TIGR03677, TIGR00432, PF13685.1, PF00832.15, TIGR00549, PF01982.11, PF00958.17, PF01864.12, PF01849.13, PF01912.13, TIGR00670, PF06026.9, PF01922.12, PF04127.10, PF05221.12, TIGR00336, TIGR01213).

### Functional annotation of genomes

An annotation pipeline was conducted to analyze genomes comparatively on the local server. Briefly speaking, putative protein-coding sequences (CDS) of 28 MAGs were identified using Prodigal v2.6.3 under the “-p single” model. Functional annotations were implemented by comparing predicted CDSs against KEGG database (Kanehisa et al. 2017) with KofamKOALA (Aramaki et al. 2020) and arCOG (Makarova et al. 2015) databases were performed. DIAMOND v2.0.11.149 (Buchfink et al. 2021) was used to implement the comparisons against the databases above, with a cutoff of 1e-5. Carbohydrate-active enzymes (CAZy) (Lombard et al. 2014) were annotated using the local version of dbcan2 (dbscan) with the parameters as: --hmm_eval 1e-5 --dia_eval 1e-5 (Zhang et al. 2018). Annotation results were kept if CAZys were identified by at least two of the three tools (HMMER, DIAMOND, and eCAMI (Xu et al. 2020)). rRNA-coding regions were determined by using barrnap v0.9 (https://github.com/tseemann/barrnap). To identify tRNA, tRNAscan-SE v2.0.9 (Lowe and Chan 2016) was deployed to all MAGs. Conserved domains of certain proteins were identified using the CD-Search tool (https://www.ncbi.nlm.nih.gov/Structure/bwrpsb/bwrpsb.cgi) on the NCBI Conserved Domain Database.

### Phylogenetic and phylogenomic analysis

286 reference genomes including 33 Parvarchaeal-associated genomes, 218 genomes from other DPANN lineages, and 35 from TACK were carefully selected from public databases for phylogenomic analyses (Supplementary Table 7). As reported in a former study, 53-concatenated-protein were selected to reconstruct phylogeny (Rinke et al. 2021). These marker sequences were identified by AMPHORA2 (Wu and Scott 2012) and aligned using MUSCLE v3.8.31 (Edgar 2004) by iterating 100 times. Poorly aligned regions were trimmed via TrimAl v1.4.rev22 (Capella-Gutierrez et al. 2009), setting the parameters as -gt 0.95 -cons 50. The phylogenetic tree was constructed using IQ-TREE v1.6.11 (Nguyen et al. 2015) with the parameter: -alrt 1000 -nt AUTO.

Reference RuBisCo sequences were selected from a previous study (Jaffe et al. 2019). These sequences were aligned with MAFFT v6.864b (Katoh and Standley 2013). Poorly aligned regions were removed using TrimAl v1.4. rev22. The unrooted phylogeny was generated using IQ-TREE with the following parameters: -alrt 1000 -bb 1000 -m JTT.

### Comparative genomics analyses

Amino acid identity (AAI) between each genome pair was determined by calculating the mean identities of orthologues, which were extracted from all reciprocal best BLAST hits (rBBHs; E-value < 1e-10). Heatmaps of AAI and amino acids usage were generated by the pheatmap package (v1.0.12) (Kolde 2019) in R. Rate smoothing of the phylogenetic tree on the heatmaps was conducted with makeChronosCalib() and chronos() functions of the ape package (v5.6-2) (Paradis and Schliep 2019). The optimal growth temperatures (OGT) were calculated using the OGT_prediction program (Sauer and Wang 2019), Tome (Li et al. 2019), and amino acids frequencies of seven amino acids (IVYWREL) (Zeldovich et al. 2007). Principle coordinate analysis (PCoA) was conducted based on the copy number of arCOGs/KOs detected in each MAG with vegan package (v2.6-2) in R (Oksanen et al. 2019). Genomic feature differences were tested by Wilcoxon signed-rank test using wilcox.test() function in stat package (v4.1.1) (Team 2016) in R. Data visualizations were conducted with ggplot2 package (v3.3.6) (Wickham 2016a).

The calculation of isoelectric point (*pI*) based on amino acid sequences of each genome was conducted by the standalone version of the isoelectric point calculator (IPC, http://isoelectric.org/) program (Kozlowski 2016) trained with the data of proteome-*pI* database (http://isoelectricpointdb.org/) (Kozlowski 2017). The proteome of each genome was further divided into “acidic” and “non-acidic” proteins according to their *pI* values. The breakpoint of the two categories was determined by the “though” of *pI* probability density function of the associated proteome of the genome, which was further used to calculate *pI* bias of the associated genome.

### Ancestral traits reconstruction

To reconstruct ancestral states for traits of interest including OGT and *pI*, we applied fastAnc() function in phytools package (v1.2-0) in R (Revell 2012). This function is based on Felsenstein’s contrasts algorithm (Felsenstein 1985) which could be used for maximum likelihood estimation (MLE) for the ancestral state. The reconstructed ancestral states were mapped with contMap() function in the same package.

## TAXONOMIC APPENDIX

### Description of ‘*Candidatus* Jingweiarchaeum’ gen. nov

‘*Candidatus* Jingweiarchaeum’ (Jing.wei.ar.chae’um. Mandarin, Jingwei, a bird from ancient Chinese mythology who was originally the daughter of an ancient Chinese ruler “Yan (Flame) Emperor” was drowned when playing alone on the seashore of the Eastern Sea. After her rebirth and becoming the “Jingwei” bird, albeit, with her tiny figure, she became determined to fill up and defeat the ocean with wood and rocks piece by piece.); N.L. neut. n. archaeum, archaeon, from Gr. adj. archaios –ê –on, ancient; N.L. neut. n. *Jingweiarchaeum*, the archaeon was named after Jingwei for her perseverance, which is similar to DPANN archaea’s strive and survival in extreme environments despite the tiny cell sizes and limited metabolic capacities.

### Description of ‘*Candidatus* Jingweiarchaeum tengchongense’ sp. nov

‘*Candidatus* Jingweiarchaeum tengchongense’ (teng.chong.gen’se. N.L. neut. adj. tengchongense, referring to Tengchong county, Yunnan Province, China, where its first genome was reconstructed).

### Description of ‘*Candidatus* Jingweiarchaeaceae’ fam. nov

‘*Candidatus* Jingweiarchaeaceae’ (Jing.wei.ar.chae.a.ce’ae. N.L. neut. n. *Jingweiarchaeum* a candidate genus; -*aceae*, ending to denote a family; N.L. fem. pl. n. *Jingweiarchaeaceae*, the *Jingweiarchaeum* candidate family).

### Description of ‘*Candidatus* Jingweiarchaeales’ ord. nov

‘*Candidatus* Jingweiarchaeales’ (Jing.wei.ar.chae.a’les. N.L. neut n. *Jingweiarchaeum* a candidate genus; -ales, ending to denote an order; N.L. fem. pl. n. *Jingweiarchaeales* the *Jingweiarchaeum* candidate order).

### Description of ‘*Candidatus* Haiyanarchaeum’ gen. nov

‘*Candidatus* Haiyanarchaeum’ (Hai.yan.ar.chae’um. Mandarin language, “Haiyan” was the husband of Jingwei that was said to be moved by her perseverance to fight with nature and join the arduous journey to defeat the ocean. N.L. neut. n. archaeum, archaeon, from Gr. adj. archaios –ê –on, ancient; N.L. neut. n. *Haiyanarchaeum*).

### Description of ‘*Candidatus* Haiyanarchaeum thermophilum’ sp. nov

*‘Candidatus* Haiyanarchaeum thermophilum’ (ther.mo’phi.lum. Gr. fem. n. therme, heat; Gr. masc. adj. philos, loving; N.L. neut. adj. thermophilum, heat-loving. Haiyanarchaeum was found in sampling sites of ultra-high temperatures and potentially, this genus has the characteristics of high estimated optimal growth temperature (OGT)).

### Description of ‘*Candidatus* Haiyanarchaeaceae’ fam. nov

‘*Candidatus* Haiyanarchaeaceae’ (Hai.yan.ar.chae.a.ce’ae. N.L. neut. n. *Haiyanarchaeum* a candidate genus; -*aceae*, ending to denote a family; N.L. fem. pl. n. *Haiyanarchaeaceae*, the *Haiyanarchaeum* candidate family).

### Description of ‘*Candidatus* Rehaiarchaeum’ gen. nov

‘*Candidatus* Rehaiarchaeum’ (Re.hai.ar.chae’um. N.L. neut. n. archaeum, an archaeon; N.L. neut.n. Rehaiarchaeum, an archaeon from R ehai National Park, Tengchong, Yunnan Province, China).

### Description of ‘*Candidatus* Rehaiarchaeum fermentans’ sp. nov

‘*Candidatus* Rehaiarchaeum fermentans’ (fer.men’tans. L. part. adj. *fermentans*, fermenting; The novel species rely on fermentation for energy production, which is different from other lineages of Parvarchaeum).

### Description of ‘*Candidatus* Parvarchaeum tengchongense’ sp. nov

‘*Candidatus* Parvarchaeum tengchongense’ (teng.chong.en’se. N.L. neut. adj. tengchongense, referring to Tengchong county. The genome was recovered from Tengchong county, Yunnan Province, China).

### Data availability

Metagenome-assembled genomes described in this study have been deposited to NCBI under the BioProject PRJNA544494: BioSample id SAMN31079384 to SAMN31079411. The data sets generated during and/or analyzed during the current study are available from the corresponding author upon reasonable request.

## Acknowledgments

We thank Guangdong Magigene Biotechnology Co., Ltd. China for the assistance in data analysis and the entire staff from Yunnan Tengchong Volcano and Spa Tourist Attraction Development Corporation for strong support. This work was financially supported by the National Natural Science Foundation of China (Nos. 91951205, GG2400000013 and 32200002). The authors are grateful to Prof. Aharon Oren (The Hebrew University of Jerusalem, Israel) for helping with the etymology.

## Author contributions

Y.Z.R., Y.X.L., Z.S.H., and W.J.L. conceived the study. Y.Z.R., Y.X.L., Y.L.Q., and Y.N.Q., performed the measurement of physicochemical parameters and DNA extraction. Y.Z.R., Y.X.L., L.Z.W., Z.S.H., Y.L.Q., J.Y.J., and Y.N.Q. performed the metagenomic analysis, genome binning, functional annotation, and evolutionary analysis. Y.Z.R., Y.X.L., L.Z.W., Z.S.H., W.J.L., and W.S.S. wrote the manuscript. All authors discussed the results and commented on the manuscript.

## Competing interests

The authors declare no competing interests.

## Supplementary Figure Legends

**Supplementary Figure 1. Phylogenetic placement of newly found genomes** with the inclusion of Huberarchaeota and based on **a)** the phylogenetic placement of all 60 MAGs in this study and **b)** the phylogenetic placement of 26 genomes after the selection of dRep with the threshold of 99% average nucleotide identity (ANI). For bootstraps in each ancestral node, values ≥ 90 were shown in purple solid circles, and values between 80 to 90 were shown in hollow circles.

**Supplementary Figure 2. Relative abundance of Jingweiarchaeales and Parvarchaeales, sampling temperatures and pH across each sampling site**. Relative abundance of the two order were shown in histograms (orange color for Jingweiarchaeales and green color for Parvarchaeles), while pH and temperatures were shown in orange and green lines respectively.

**Supplementary Figure 3. A maximum likelihood phylogenetic tree of the large subunit of ribulose-1**,**5-bisphosphate (RuBP) carboxylase–oxygenase (RuBisCO)**. For bootstraps in each ancestral node, values ≥ 75 were shown in grey solid circles. The names of MAGs in this study were labeled in brown.

**Supplementary Figure 4. Other genomic features statistics of all MAGs in this study besides Fig. 3**. Differences of **a)** predicted genome size (48 SCGs), **b)** predicted genome size (CheckM2), **c)** predicted genome size (CheckM1), **d)** average gene length, **e)** coding density, **f)** ratio of overlapping genes, **g)** GC content, **h)** average isoelectric point (*pI*), **i)** optimal growth temperatures (OGT) predicted with the approach of Wang^103^ et al., and **j)** OGT predicted by Tome^104^ were visualized using ggplot2 package (v3.3.6)^105^. Different alphabets show the significant differences between groups with P <0.005.

**Supplementary Figure 5. Principle coordinate analysis (PCoA) of Jingweiarchaeum, Haiyanarchaeum, Rehaiarchaeum, Acidifodinimicrobium, and Parvarchaeum**. Genomes were plotted with the main axes of PCoA according to the annotation of KEGG and arCOG databases. Squares represent genomes of Jingweiarchaeales and triangles represent genomes of Parcarchaeales. While the colors of tshapes represent the associate sampling environments in which MAGs recovered from hot spring were shown in green and MAGs recovered from acid mine drainages (AMD) were shown in blue.

**Supplementary Figure 6. Average amino acid identity (AAI) heatmap**. Hierarchical clustering heatmap based on AAI. The dendrogram shown on the left was extracted from the maximum likelihood tree (Fig. 1) to illustrate the phylogenetic relationship of Jingweiarchaeum, Haiyanarchaeum, Rehaiarchaeum, Acidifodinimicrobium, and Parvarchaeum.

**Supplementary Figure 7. Heatmap of amino acids usage of all MAGs in this study**. The lateral axis represents 20 amino acids while the vertical axis represents MAGs in this study, with the order of phylogenic placement. The blue to red colors show the accumulation of each amino acid for each MAG after the Z-score transformation.

**Supplementary Figures 8-13. Isoelectric point (*pI*) and molecular weight of the predicted proteins from each MAG in this study**. The vertical axis represents molecular weight, and the lateral axis represents the *pI* of the predicted protein from each MAG.

**Supplementary Figure 14. A maximum likelihood phylogenetic tree constructed for reverse gyrase**. For bootstraps in each ancestral node, values ≥ 75 were shown in grey solid circles. The reverse gyrase sequence presents in the MAG JZ-1_201705_bins_113 that is affiliated with Haiyanarchaeum is labeled in brown.

**Supplementary Figure 15. A maximum likelihood phylogenetic tree constructed for superoxide dismutase (SODA)**. For bootstraps in each ancestral node, values ≥ 75 were shown in grey solid circles. The names of MAGs in this study were labeled brown. Sequences of the three newly discovered genara, Rehaiarchaeaum, Acidifodinimicrobium, and Parvarchaeum, were labeled with purple, blue and orange hexagons, respectively.

**Supplementary Figure 16. Linear regressions of genome sizes of Jingweiarchaeum and Haiyanarchaeum versus sampling site temperatures and OGTs**. Green circles represent the genomic features of Jingweiarchaeum while red circles represent the genomic features of Haiyanarchaeum.

**Supplementary Figure 17. A maximum likelihood phylogenetic tree constructed for AhpC/TSA family peroxiredoxin**. For bootstraps in each ancestral node, values ≥ 75 were shown in grey solid circles. Sequences of the names of MAGs in this study were labeled brown. Three newly discovered genara, Rehaiarchaeaum, Acidifodinimicrobium, and Parvarchaeum, were labeled with purple, blue, and orange hexagons, respectively.

**Supplementary Figure 18. A maximum likelihood phylogenetic tree constructed for peroxiredoxin**. For bootstraps in each ancestral node, values ≥ 75 were shown in grey solid circles. The names of MAGs in this study were labeled brown. Sequences of three newly discovered genera, Rehaiarchaeaum, Acidifodinimicrobium, and Parvarchaeum, were labeled with purple, blue, and orange hexagons, respectively.

## Supplementary Table Legends

**Supplementary Table 1. General genome statistics, genome taxonomy, and sampling/predicted physiological characteristics of each organism represented by the associated genome**.

**Supplementary Table 2. Geochemical properties of sampling sites for all metagenomes**

**Supplementary Table 3. Occurrences of 48 DPANN-specific single-copy genes (SCGs) present in 60 Parvarchaeales-related genomes**

**Supplementary Table 4. Occurrences of 149 CheckM1 archaeal markers in 60 Parvarchaeales-related genomes**

**Supplementary Table 5. Occurrences of 53 GTDB archaeal markers in all genomes selected for phylogenetic analyses**

**Supplementary Table 6. Summary of metabolic potentials of 58 Jingweiarchaeales and Parvarchaeales genomes**

**Supplementary Table 7. List of reconstructed ancestral traits including OGT (IVYWREL), and OGT (Tome), *pI* bias, and average *pI***.

## Additional Data

**Additional Data 1: Annotation of all MAGs to KofamKOALA databases.**

**Additional Data 2: Annotation of all MAGs to arCOG databases**.

## Reference

Alneberg, J., Bjarnason, B.S., Bruijn I.D., Schirmer, M., Quick, J, Ijaz, U.Z., Lahti, L., Loman, N.J., Andersson, A.F., Quince, C. (2014). Binning metagenomic contigs by coverage and composition. Nature methods 11, 1144–1146.

Aramaki, T., Blanc-Mathieu, R., Endo, H., Ohkubo, K., Kanehisa, M, Goto, S., and Ogata, H. (2020). KofamKOALA: KEGG Ortholog assignment based on profile HMM and adaptive score threshold. Bioinformatics 36, 2251–2252.

Baker, B.J., Comolli, L.R., Dick, G.R., Hauser, L.J., Hyatt, D., Dill, B.D., Land, M.L., VerBerkmoes, N.C., Robert L., Banfield, J.F. (2010). Enigmatic, ultrasmall, uncultivated Archaea. Proceedings of the National Academy of Sciences 107, 8806–8811.

Baker, B.J., De Anda, V., Seitz, K.W., Dombrowski, N., Santoro, A.E., Lloyd, K.G. (2020). Diversity, ecology and evolution of Archaea. Nature Microbiology 5, 887–900.

Baker, B.J., Tyson, G.W., Webb, R.I., Flanagan, J., Hugenholtz, P, Allen, E.E., and Banfield, J.F. (2006). Lineages of acidophilic archaea revealed by community genomic analysis. Science 314, 1933–1935.

Baker-Austin, C., Dopson, M. (2007). Life in acid: pH homeostasis in acidophiles. Trends in Microbiology 15, 165–171.

Bankevich, A., Nurk, S., Antipov, D., Gurevich, A.A., Dvorkin, M., Kulikov, A.S., Lesin, V.M., Nikolenko, S.I., Pham, S., Prjibelski, A.D., et al. (2012). SPAdes: a new genome assembly algorithm and its applications to single-cell sequencing. Journal of Computational Biology 19, 455–477.

Beam, C.E., Saveson, C.J., Lovett, S.T. (2002). Role for radA/sms in recombination intermediate processing in Escherichia coli. Journal of Bacteriology 184, 6836–6844.

Beam, J.P., Becraft, E.D., Brown, J.M., Schulz, F., Jarett, J.K., Bezuidt, O., Poulton, N.J., Clark, K, Dunfield, P.F., Ravin, N.V., et al. (2020). Ancestral absence of electron transport chains in Patescibacteria and DPANN. Frontiers in Microbiology 11, 1848.

Bergsten, J. (2005). A review of long-branch attraction. Cladistics 21, 163–193.

Borisov, V.B., Gennis, R.B., Hemp, J., and Verkhovsky, M.I. (2011). The cytochrome bd respiratory oxygen reductases. Biochimica et Biophysica Acta (BBA)-Bioenergetics 1807, no. 11 (2011): 1398–1413.

Buchfink, B., Reuter, K., Drost, H.-G. (2021). Sensitive protein alignments at tree-of-life scale using DIAMOND. Nature Methods 18, 366–368.

Capella-Gutierrez, S., Silla-Martinez, J.M., Gabaldon, T. (2009). trimAl: a tool for automated alignment trimming in large-scale phylogenetic analyses. Bioinformatics 25, 1972–1973.

Castelle, C.J., Brown, C.T., Anantharaman, K., Probst, A.J., Huang, R.H., Banfield, J.F. (2018). Biosynthetic capacity, metabolic variety and unusual biology in the CPR and DPANN radiations. Nature Reviews Microbiology 16, 629–645.

Castelle, C.J., Wrighton, K.C., Thomas, B.C., Hug, L.A., Brown, C.T., Wilkins, M.J., Frischkorn, K.R., Tringe, S.G., Singh, A., et al. (2015). Genomic expansion of domain archaea highlights roles for organisms from new phyla in anaerobic carbon cycling. Current Biology 25, 690–701.

Chaumeil, P.-A., Mussig, A.J., Hugenholtz, P., Parks, D.H. (2019). GTDB-Tk: a toolkit to classify genomes with the Genome Taxonomy Database. Bioinformatics 36, 1925–1927

Chen, L.-X., Méndez-García, C., Dombrowski, N., Servín-Garcidueñas, L.E., Eloe-Fadrosh, E.A., Fang, B.-Z., Luo, Z.H., Tan, S., Zhi, X.Y., Hua Z.S. et al. (2018). Metabolic versatility of small archaea Micrarchaeota and Parvarchaeota. The ISME journal 12, 756–775.

Chklovski, A., Parks, D.H., Woodcroft, B.J., Tyson, G.W. (2022). CheckM2: a rapid, scalable and accurate tool for assessing microbial genome quality using machine learning. bioRxiv

Comolli, L.R., Banfield, J.F. (2014). Inter-species interconnections in acid mine drainage microbial communities. Frontiers in Microbiology 5, 367.

Dietl, A., Ferousi, C., Maalcke, W.J., Menzel, A., de Vries, S., Keltjens, J.T., Jetten, M.S.M, Kartal, B., and Barends, T.R.M. (2015). The inner workings of the hydrazine synthase multiprotein complex. Nature 527, 394–397.

Dombrowski, N., Williams, T.A., Sun, J., Woodcroft, B.J., Lee, J.-H., Minh, B.Q., Rinke, C., and Spang, A. (2020). Undinarchaeota illuminate DPANN phylogeny and the impact of gene transfer on archaeal evolution. Nature Communications 11, 3939.

Edgar, R.C. (2004). MUSCLE: multiple sequence alignment with high accuracy and high throughput. Nucleic Acids Research 32, 1792–1797.

Feller, G. (2013). Psychrophilic enzymes: from folding to function and biotechnology. Scientifica 2013, 512840.

Felsenstein, J. (1985). Phylogenies and the comparative method. The American Naturalist 125, 1–15.

Fukui, T., Atomi, H., Kanai, T., Matsumi, R., Fujiwara, S., Imanaka, T. (2005). Complete genome sequence of the hyperthermophilic archaeon Thermococcus kodakaraensis KOD1 and comparison with Pyrococcus genomes. Genome Research 15, 352–363.

Galtier, N., Lobry, J.R. (1997). Relationships between genomic G+C Content, RNA secondary structures, and optimal growth temperature in prokaryotes. Journal of Molecular Evolution 44, 632–636.

Glasemacher, J., Bock, A.-K., Schmid, R., Schönheit, P. (1997). Purification and properties of acetyl-CoA synthetase (ADP-forming), an archaeal enzyme of acetate formation and ATP synthesis, from the hyperthermophile Pyrococcus furiosus. European Journal of Biochemistry 244, 561–567.

Groussin, M., Boussau, B., Charles, S., Blanquart, S., Gouy, M. (2013). The molecular signal for the adaptation to cold temperature during early life on Earth. Biology Letters 9, 20130608.

Groussin, M., Gouy, M. (2011). Adaptation to environmental temperature is a major determinant of molecular evolutionary rates in archaea. Molecular Biology and Evolution 28, 2661–2674.

He, Christine, Ray Keren, Michael L. Whittaker, Ibrahim F. Farag, Jennifer A. Doudna, Jamie HD Cate, and Jillian F. Banfield. (2021). Genome-resolved metagenomics reveals site-specific diversity of episymbiotic CPR bacteria and DPANN archaea in groundwater ecosystems. Nature Microbiology 6, 354–365.

Heine, M., Chandra, S.B.C. (2009). The linkage between reverse gyrase and hyperthermophiles: A review of their invariable association. The Journal of Microbiology 47, 229–234.

Hua, Z.-S., Qu, Y.-N., Zhu, Q., Zhou, E.-M., Qi, Y.-L., Yin, Y.-R., Rao Y.Z., Tian, Y., Li, Y.X., Liu, L., et al. (2018). Genomic inference of the metabolism and evolution of the archaeal phylum Aigarchaeota. Nature Communications 9, 2832.

Huber, H., Hohn, M.J., Rachel, R., Fuchs, T., Wimmer, V.C., Stetter, K.O. (2002). A new phylum of Archaea represented by a nanosized hyperthermophilic symbiont. Nature 417, 63–67.

Jaffe, A.L., Castelle, C.J., Dupont, C.L., Banfield, J.F. (2019). Lateral gene transfer shapes the distribution of RuBisCO among candidate phyla radiation bacteria and DPANN Archaea. Molecular Biology and Evolution 36, 435–446.

Kampmann, M., Stock, D. (2004). Reverse gyrase has heat-protective DNA chaperone activity independent of supercoiling. Nucleic Acids Research 32, 3537–3545.

Kanehisa, M., Furumichi, M., Tanabe, M., Sato, Y., Morishima, K. (2017). KEGG: new perspectives on genomes, pathways, diseases and drugs. Nucleic Acids Research 45, D353–D361.

Kang, D.D., Froula, J., Egan, R., Wang, Z. MetaBAT: Metagenome binning based on abundance and tetranucleotide frequency. (2014). eScholarship 4.

Kartal, B., Maalcke, W.J., de Almeida, N.M., Cirpus, I., Gloerich, J., Geerts, W., Huub JM Op den Camp, Harhangi, H.R., Janssen-Megens, E.M., Francoijs, K., et al. (2011). Molecular mechanism of anaerobic ammonium oxidation. Nature 479, 127–130.

Katoh, K., Standley, D.M. (2013). MAFFT Multiple Sequence Alignment Software Version 7: Improvements in Performance and Usability. Molecular Biology and Evolution 30, 772–780.

Kiraga, J., Mackiewicz, P., Mackiewicz, D., Kowalczuk, M., Biecek, P., Polak, N., et al. (2007). The relationships between the isoelectric point and: length of proteins, taxonomy and ecology of organisms. BMC Genomics 8, 163.

Knoll, A.H., Nowak, M.A. (2017). The timetable of evolution. (2017). Science Advances 3, no. 5: e1603076.

Kolde, R. (2019). pheatmap: Pretty Heatmaps. R package version 1.0. 12

Kozlowski, L.P. (2016). IPC – Isoelectric Point Calculator. Biology Direct 11, 55.

Kozlowski, L.P. (2017). Proteome-pI: proteome isoelectric point database. Nucleic Acids Research 45, D1112–D1116.

Lewin, A., Wentzel, A., Valla, S. (2013). Metagenomics of microbial life in extreme temperature environments. Current Opinion in Biotechnology 24, 516–525.

Li, G., Rabe, K.S., Nielsen, J., Engqvist, M.K.M. (2019). Machine learning applied to predicting microorganism growth temperatures and enzyme catalytic optima. ACS synthetic biology 8, 1411–1420.

Li, Y.F., Costello, J.C., Holloway, A.K., Hahn, M.W. (2008). “Reverse Ecology” and the Power of Population Genomics. Evolution: International Journal of Organic Evolution 62, 2984–2994.

Li, Y.-X., Rao, Y.-Z., Qi, Y.-L., Qu, Y.-N., Chen, Y.-T., Jiao, J.-Y., et al. (2021). Deciphering Symbiotic Interactions of “Candidatus Aenigmarchaeota” with Inferred Horizontal Gene Transfers and Co-occurrence Networks. mSystems 6, e00606–21.

Liu, X., Li, M., Castelle, C.J., Probst, A.J., Zhou, Z., Pan, J, Liu, Y., Banfield, J.F., and Gu, J.D. (2018). Insights into the ecology, evolution, and metabolism of the widespread Woesearchaeotal lineages. Microbiome 6, 102.

Lombard, V., Golaconda Ramulu, H., Drula, E., Coutinho, P.M., Henrissat, B. (2014). The carbohydrate-active enzymes database (CAZy) in 2013. Nucleic acids research 42, D490–D495.

Lowe, T.M., Chan, P.P. (2016). tRNAscan-SE On-line: integrating search and context for analysis of transfer RNA genes. Nucleic Acids Research 44, W54–W57.

Lulchev, P., Klostermeier, D. (2014). Reverse gyrase—recent advances and current mechanistic understanding of positive DNA supercoiling. Nucleic Acids Research 42, 8200–8213.

Luo, Z.-H., Li, Q., Lai, Y., Chen, H., Liao, B., Huang, L. (2020). Diversity and genomic characterization of a novel Parvarchaeota family in acid mine drainage sediments. Frontiers in Microbiology 11, 612257.

Makarova, K., Wolf, Y., Koonin, E. (2015). Archaeal Clusters of Orthologous Genes (arCOGs): an update and application for analysis of shared features between Thermococcales, methanococcales, and methanobacteriales. Life 5, 818–840.

Makarova, K.S. (2016). Diversity and evolution of type IV pili systems in archaea. Frontiers in Microbiology 7, 16.

Najmanovich, R., Kuttner, J., Sobolev, V., Edelman, M. (2000). Side-chain flexibility in proteins upon ligand binding. Proteins: Structure, Function, and Bioinformatics 39, 261–268.

Nguyen, L.-T., Schmidt, H.A., von Haeseler, A., Minh, B.Q. (2015). IQ-TREE: a fast and effective stochastic algorithm for estimating maximum-likelihood phylogenies. Molecular Biology and Evolution 32, 268–274.

Oksanen, J., Blanchet, F.G., Friendly, M., Kindt, R., Legendre, P., McGlinn, D., P. R. Minchin et al. (2019). vegan: Community Ecology Package. R package (Version 2.5–4)

Olm, M.R., Brown, C.T., Brooks, B., Banfield, J.F. (2017). dRep: a tool for fast and accurate genomic comparisons that enables improved genome recovery from metagenomes through dereplication. The ISME journal 11, 2864–2868.

Paradis, E., Schliep, K. (2019). ape 5.0: an environment for modern phylogenetics and evolutionary analyses in R. Bioinformatics 35, 526–528.

Parks, D.H., Imelfort, M., Skennerton, C.T., Hugenholtz, P., Tyson, G.W. (2015). CheckM: assessing the quality of microbial genomes recovered from isolates, single cells, and metagenomes. Genome Research 25, 1043–1055.

Parks, D.H., Rinke, C., Chuvochina, M., Chaumeil, P.-A., Woodcroft, B.J., Evans, P.N., et al. (2017). Recovery of nearly 8,000 metagenome-assembled genomes substantially expands the tree of life. Nature Microbiology 2, 1533–1542.

Poole, R.K., Hill, S. (1997). Respiratory protection of nitrogenase activity in Azotobacter vinelandii—roles of the terminal oxidases. Bioscience Reports 17, 303–317.

Probst, A.J., Ladd, B., Jarett, J.K., Geller-McGrath, D.E., Sieber, C.M.K., Emerson, J.B., Anantharaman, K., Thomas, B.C., Malmstrom, R.R., et al. (2018). Differential depth distribution of microbial function and putative symbionts through sediment-hosted aquifers in the deep terrestrial subsurface. Nature Microbiology 3, 328–336.

Ram, R.J., VerBerkmoes, N.C., Thelen, M.P., Tyson, G.W., Baker, B.J., Blake, R.C., Shah, M., Hettich, R.L., Banfield, J.F. (2005). Community Proteomics of a Natural Microbial Biofilm. Science 308, 1915–1920.

Reeves, R.E., Warren, L.G., Susskind, B., Lo, H.S. (1977). An energy-conserving pyruvate-to-acetate pathway in Entamoeba histolytica. Pyruvate synthase and a new acetate thiokinase. Journal of Biological Chemistry 252, 726–731.

Revell, L.J. (2012). phytools: an R package for phylogenetic comparative biology (and other things). Methods in Ecology and Evolution, 217–223.

Rinke, C., Chuvochina, M., Mussig, A.J., Chaumeil, P.-A., Davín, A.A., Waite, D.W., et al. (2021). A standardized archaeal taxonomy for the Genome Taxonomy Database. Nature Microbiology 6, 946–959.

Rinke, C., Schwientek, P., Sczyrba, A., Ivanova, N.N., Anderson, I.J., Cheng, J.-F., et al. (2013). Insights into the phylogeny and coding potential of microbial dark matter. Nature 499, 431–437.

Sabath, N., Ferrada, E., Barve, A., Wagner, A. (2013). Growth temperature and genome size in bacteria are negatively correlated, suggesting genomic streamlining during thermal adaptation. Genome Biology and Evolution 5, 966–977.

Sakaguchi, M., Matsuzaki, M., Niimiya, K., Seino, J., Sugahara, Y., Kawakita, M. (2007). Role of proline residues in conferring thermostability on aqualysin I. The Journal of Biochemistry 141, 213–220.

Sakai, H.D., Nur, N., Kato, S., Yuki, M., Shimizu, M., Itoh, T., et al. (2022). Insight into the symbiotic lifestyle of DPANN archaea revealed by cultivation and genome analyses. Proceedings of the National Academy of Sciences 119, e2115449119.

Sato, T., Atomi, H., Imanaka, T. (2007). Archaeal type III rubiscos function in a pathway for amp metabolism. Science 315, 1003–1006.

Sauer, D.B., Wang, D.N. (2019). Predicting the optimal growth temperatures of prokaryotes using only genome derived features. Bioinformatics 35, 3224–3231.

Say, R.F., Fuchs, G. (2010). Fructose 1,6-bisphosphate aldolase/phosphatase may be an ancestral gluconeogenic enzyme. Nature 464, 1077–1081.

Shu, W.S., Huang, L.N. (2022). Microbial diversity in extreme environments. Nature Reviews Microbiology 20, 219–235.

Siddiqui, K.S., Poljak, A., Guilhaus, M., De Francisci, D., Curmi, P.M.G., Feller, G., D’Amico, S., Gerday, C., Uversky, V.N., and Cavicchioli, R. (2006). Role of lysine versus arginine in enzyme cold-adaptation: Modifying lysine to homo-arginine stabilizes the cold-adapted α-amylase from Pseudoalteramonas haloplanktis. Proteins: Structure, Function, and Bioinformatics 64, 486–501.

Sieber, C.M.K., Probst, A.J., Sharrar, A., Thomas, B.C., Hess, M., Tringe, S.G., and Banfield, J.F. (2018). Recovery of genomes from metagenomes via a dereplication, aggregation and scoring strategy. Nature Microbiology 3, 836–843.

Stetter, K.O. (2006). Hyperthermophiles in the history of life. Philosophical Transactions of the Royal Society B: Biological Sciences 361, 1837–1843.

R Core Team. (2016). R: A language and environment for statistical computing [Computer software manual]. Vienna, Austria.

Tyson, G.W., Chapman, J., Hugenholtz, P., Allen, E.E., Ram, R.J., Richardson, P.M., Victor V. Solovyev, Edward M. Rubin, Daniel S. Rokhsar, and Jillian F. Banfield. (2004). Community structure and metabolism through reconstruction of microbial genomes from the environment. Nature 428, 37–43.

Vázquez-Campos, X., Kinsela, A.S., Bligh, M.W., Payne, T.E., Wilkins, M.R., Waite, T.D. (2021). Genomic insights into the archaea inhabiting an australian radioactive legacy site. Frontiers in Microbiology 12, 3069.

Venditti, C., Meade, A., Pagel, M. (2006). Detecting the node-density artifact in phylogeny reconstruction. Systematic Biology 55, 637–643.

White, A.D., Keefe, A.J., Ella-Menye, J.-R., Nowinski, A.K., Shao, Q., Pfaendtner, J., and Jiang, S. (2013). Free energy of solvated salt bridges: a simulation and experimental study. The Journal of Physical Chemistry B 24, 7254–7259

Wickham, H. (2016a). ggplot2: Elegant Graphics for Data Analysis. 2nd ed. (2016). Springer.

Williams, T.A., Cox, C.J., Foster, P.G., Szöllősi, G.J., Embley, T.M. (2020). Phylogenomics provides robust support for a two-domains tree of life. Nature Ecology & Evolution 4, 138–147.

Williams, T.A., Szöllősi, G.J., Spang, A., Foster, P.G., Heaps, S.E., Boussau, B., Thijs JG Ettema, and T. Martin Embley (2017). Integrative modeling of gene and genome evolution roots the archaeal tree of life. Proceedings of the National Academy of Sciences 114, E4602–E4611.

Woese, C.R., Fox, G.E. (1977). Phylogenetic structure of the prokaryotic domain: The primary kingdoms. Proceedings of the National Academy of Sciences 74, 5088–5090.

van Wolferen, M., Orell, A., Albers, S.-V. (2018). Archaeal biofilm formation. Nature Reviews Microbiology 16, 699–713.

Wrighton, K.C., Castelle, C.J., Varaljay, V.A., Satagopan, S., Brown, C.T., Wilkins, M.J., et al. (2016). RubisCO of a nucleoside pathway known from Archaea is found in diverse uncultivated phyla in bacteria. The ISME journal 10, 2702–2714.

Wu, M., Scott, A.J. (2012). Phylogenomic analysis of bacterial and archaeal sequences with AMPHORA2. Bioinformatics 28, 1033–1034.

Wu, Y.-W., Simmons, B.A., Singer, S.W. (2016). MaxBin 2.0: an automated binning algorithm to recover genomes from multiple metagenomic datasets. Bioinformatics 32, 605–607.

Xu, J., Zhang, H., Zheng, J., Dovoedo, P., Yin, Y. (2020). eCAMI: simultaneous classification and motif identification for enzyme annotation. Bioinformatics 36, 2068–2075.

Youssef, N.H., Rinke, C., Stepanauskas, R., Farag, I., Woyke, T., Elshahed, M.S. (2015). Insights into the metabolism, lifestyle and putative evolutionary history of the novel archaeal phylum ‘Diapherotrites’. The ISME journal 9, 447–460.

Zeghouf, M., Fontecave, M., Covès, J. (2000). A simplifed functional version of the escherichia coli sulfite reductase. Journal of Biological Chemistry 275, 37651–37656.

Zeldovich, K.B., Berezovsky, I.N., Shakhnovich, E.I. (2007). Protein and dna sequence determinants of thermophilic adaptation. PLoS Computational Biology 3, e5.

Zhang, H., Yohe, T., Huang, L., Entwistle, S., Wu, P., Yang, Z., Busk, P.K., Xu, Y., and Yin, Y. (2018). dbCAN2: a meta server for automated carbohydrate-active enzyme annotation. Nucleic Acids Research 46, W95–W101.

Zhaxybayeva, O., Swithers, K.S., Lapierre, P., Fournier, G.P., Bickhart, D.M., DeBoy, R.T., Karen E.N, Nesbøe, C.L., Doolittle, W.F, Gogarten, J.P., et al. (2009). On the chimeric nature, thermophilic origin, and phylogenetic placement of the Thermotogales. Proceedings of the National Academy of Sciences, 106, 5865–5870.

